# Upgrading Voxel-wise Encoding Model via Integrated Integration over Features and Brain Networks

**DOI:** 10.1101/2022.11.06.515387

**Authors:** Yuanning Li, Huzheng Yang, Shi Gu

**Author notes:** Correspondence to: Shi Gu and Yuanning Li. These authors contributed equally (Yuanning Li, Huzheng Yang).

## Abstract

A central goal of cognitive neuroscience is to build computational models that predict and explain neural responses to sensory inputs in the cortex. Recent studies attempt to borrow the representation power of deep neural networks (DNN) to predict the brain response and suggest a correspondence between artificial and biological neural networks in their feature representations. However, each DNN instance is often specified for certain computer vision tasks which may not lead to optimal brain correspondence. On the other hand, these voxel-wise encoding models focus on predicting single voxels independently, while brain activity often demonstrates rich and dynamic structures at the population and network levels during cognitive tasks. These two important properties suggest that we can improve the prevalent voxel-wise encoding models by integrating features from DNN models and by integrating cortical network information into the models. In this work, we propose a new unified framework that addresses these two aspects through DNN feature-level ensemble learning and brain atlas-level model integration. Our proposed approach leads to superior performance over previous DNN-based encoding models in predicting whole-brain neural activity during naturalistic video perception. Furthermore, our unified framework also facilitates the investigation of the brain’s neural representation mechanism by accurately predicting the neural response corresponding to complex visual concepts.

## Introduction

A central goal of computational cognitive neuroscience is to build models that explain how the brain perceives sensory information^1^. An ideal computational model of sensory perception would be able to both perform the sensory perception task behaviorally and explain the underlying neural basis during the perception process^2–6^. This implies two critical goals: to model and predict neural activity in the brain with high accuracy and to achieve human-level performance behaviorally. Previous efforts diverge along these two routines. Most studies in visual and auditory neuroscience focus on analyzing how different levels of sensory information are represented in the cortical network and link these neural coding to perceptual behavior.^4,7–15^ These hypothesis-driven works succeeded in interpreting neural coding and identifying the neural basis of behavioral properties. However, due to the limitation of linear models and the ad-hoc choices of features used in these models, these hypothesis-driven methods often fall short in predicting neural activity with high accuracy. Furthermore, these empirical results cannot be directly turned into computational agents that perform such perception tasks thus lack high-level behavioral descriptions. On the other hand, cognitive models, particularly connectionist models, are designed to mimic human sensory perceptual behavior and perform the same tasks as humans.^16,17^ It is not until the surge of deep neural networks over the past decade that these models finally approach and surpass the human level in many sensory cognition tasks.^18,19^ As opposite to the neural coding studies, these artificial neural network (DNN) models excel in computational tasks, but it remains unclear whether and to what extent they reflect the same underlying representation and computations as the neural system.

The recent advance in DNN models inspires new efforts that combine computational models with neural coding models.^5,20–24^ Specifically, these powerful pretrained networks are employed to build unit/voxel-wise prediction models in the cortex. These models fit an encoder from the external stimulus to the brain signal and allow for the investigation of representation and computations in large-scale neural circuits through the correlations between artificial neural layers and brain regions. These DNN models have already been optimized for performing corresponding cognitive tasks. As a prediction model, the main goal is to achieve high neural prediction accuracy in order to facilitate further analyses of the underlying coding and computation mechanisms.^2,6^

Previous studies using voxel-wise encoding models have shown that, compared to theory-driven heuristic models, DNN models can predict neural responses with regard to static images and sounds in different ROIs within sensory cortex with higher accuracy.^5,21,22,25^ Some recent studies have also demonstrated that these approaches can be extended to naturalistic stimuli, such as movies and speech.^23,24,26,27^ However, two important challenges have limited the prediction performance of these models. First, the brain is an interconnected network with different areas dynamically reconfigured and involved in different modules during cognitive tasks,^28–32^ while the prevalent voxel-wise encoding models treat each voxel static and independently. Second, by using pretrained task optimized DNN models, it is often assumed that there is a single optimal set of representation features aligned with a specific neural population along the network hierarchy.^6,22^ However, the feature representations are mainly driven by training objectives and enforcing a one-to-one correspondence may not be optimal. These two factors have significantly limited the performance of the current DNN-based models. Even the state-of-the-art DNN-based models can only explain ∼50% of the total variance driven by the input stimuli.^6^ Therefore, pushing the model prediction performance towards the upper limit is an urgent demand for such prediction models.

To get high encoding prediction accuracy via addressing these two issues, we focus on two sides of the encoding models. On the targeting neural activity side, it is often overlooked in the previous studies that both the stimulus-driven and the spontaneous parts of the neural activity show strong correlating structure at local and network levels.^33–36^ Thus we ask if we could incorporate correlated activity into the model by harnessing local and network local level structures in the neural activity to facilitate accurate neural encoding prediction. On the stimulus side, existing literature usually extracts feature representations from the stimuli by picking the optimal feature representation from a candidate model pool using model-selection procedures.^21,25^ However, the brain is a linked system where stimuli usually activate a broad network of cortical areas across the whole brain^9,37^, suggesting that the representation may be an integration of multi-level features rather than driven by a dominating mode. Moreover, an artificial neural network is not designed for replicating the brain topology thus different levels of feature extraction within the same model may also align to different neural populations.^38^ Thus we ask if we can push the capability of the encoding model towards the ceiling by enriching the feature representations to an integration on multiple levels over multiple regions in modeling the neural responses to naturalistic stimuli.

Following this prediction-center principle, we identify three pairs of principles in neuroscience that could benefit the prediction from the machine learning perspective and validate the efficacy based on three levels of corresponding hypotheses. First, the neural activity of the brain is reflected in functional modules that are related but not overlapped with the underlying anatomy. Voxels that are not clustered spatially may also correlated through functional networks and shared both stimulus-driven and non-stimulus endogenous activity.^35^ Thus we hypothesize that the function-induced cluster-based encoding model provides complementary prediction power to the anatomy-induced model. Second, a brain region may participate in multiple perception processes that could be better captured by different computational models.^39^ Thus we hypothesize that integrating stimulus-derived features from different processing levels within each model will improve neural encoding accuracy. Thirdly, a brain region may reconfigure its role across multiple perception processes reflected in the form of different modularized structures.^31^ Thus we hypothesize that there exists heterogeneity in model performance across different ways of ROI clustering, and integrating these different atlases further improves model performance.

Moreover, we demonstrate the efficacy of the prediction-centered model from two applying views. Since the encoding weights identify an artificial neural network, we show that it serves as a novel metric that reveals functional organizations of voxels that deviate from the pure anatomically defined ROIs. Further, based on the representation similarity scores, we show that our more accurate prediction model actually results in a more similar representation with the brain regarding visual motion. Our approach promotes insight into why we should focus on prediction in building future encoding models.

## Results

In this study, brain activity was recorded using functional magnetic resonance imaging (fMRI) when 10 subjects passively viewed 1102 naturalistic video clips. We focus on predicting the brain response from the corresponding video stimuli.^40^ We adopt the general voxel-wise neural encoding framework that has been widely used in the literature.^41–43^ In particular, DNN models are used to extract feature representations from each individual video stimulus. Another multilayer perceptron (MLP) network is trained to predict brain activation in each individual voxel regarding each stimulus, using the extracted features from the DNN models.

To do this, we developed an iterative integration approach. As demonstrated in Figure 1, our model consists of two parts of integrations: the feature-level integration and the atlas-level integration. First, features of the input stimuli were extracted via feature-level integration that ensembles features from different layers of DNN models under multiple optimization parameters (Fig. 1a). Second, atlas-level integration was performed to combine encoding models based on multiple functional and anatomical atlases (Fig. 1d). Different functional atlases were constructed based on task-optimized parcellations using encoding model weights from voxel-wise encoders (Fig. 1b). These functional atlases grouped voxels with similar representation properties together (Fig. 1c). We demonstrate the two parts of integrations and evaluate the performance of the overall model in the following sections.

**Figure 1:**
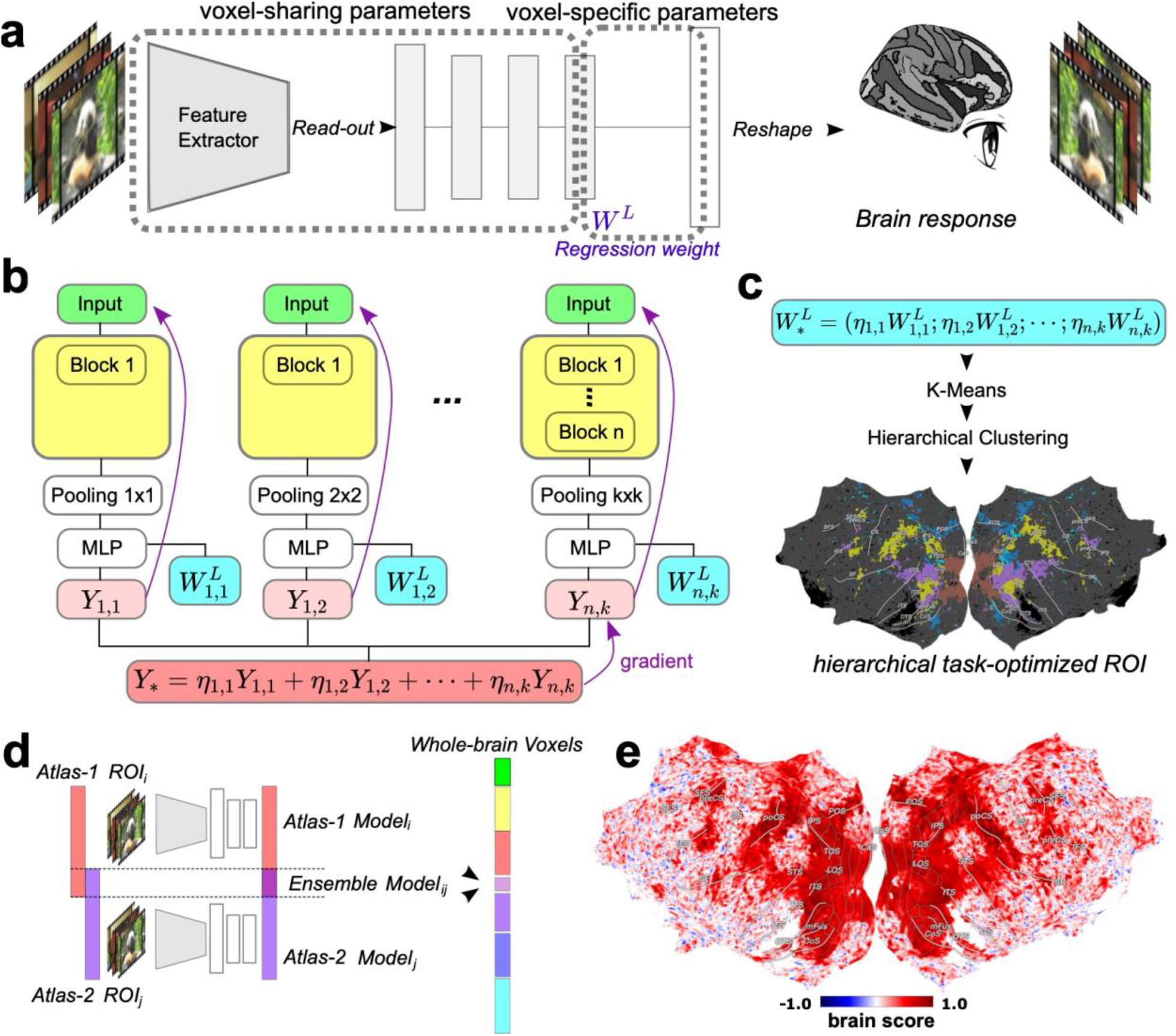
Overview of the feature-level and atlas-level integration framework. a) Overview of voxel-wise encoding model framework. We feed input videos to a pre-trained deep neural network feature extractor and take intermediate layers to a multiple-layer perceptron to predict voxel-wise whole brain response captured by fMRI. The whole model is trained end-to-end with all parameters tunable. The last layer, with voxel activations as output, can be interpreted as linear regression with weights denoted as *W*^*L*^. All the voxels share parameters except for the last linear regression. b) Overview of the feature level integration: we trained models separately while taking different intermediate layers and read-out pooling sizes, denoted as *Y*. Then we optimized an offline linear combination of their outputs with the linear weights denoted as *η*. The arrows indicate the gradient flow, and there is no gradient from the combined output to the input video. c) Functional clustering based on voxel-wise encoding weights: regression weights *W*^*L*^ are weighted by the linear combination *η*. The concatenated regression weights *W*^\**L*^ are then used as voxel embeddings for clustering. d) Atlas-level integration: each model is trained with voxels from the same ROI as output, while each atlas contains several ROIs. On different atlases, we combined the model outputs on their ROI-intersection (overlap of red and purple bars). e) Best model prediction score were plotted on the whole cortical surface, normalized to noise-ceiling.

### Feature-level integration

The prevalent practice for training a DNN-based voxel-wise encoding model depends on the strategy of choosing the best feature space with the highest prediction score,^6^ or concatenating features from multiple intermediate layers.^24^ We challenge these strategies both from neuroscience and deep neural network perspectives. Instead of these rather heuristic feature-selection strategies, we propose a systematic way of feature-integration via ensemble learning. On the one hand, there may not exist a one-to-one matching between the DNN feature layers and different neural populations, and one specific neural population may be involved in multiple different levels of information processing spanning over a set of features across the DNN hierarchy.^38^ On the other hand, the convergence speed varies when using intermediate layers and pooling sizes. For example, STS prediction model using high-level DNN features converges two times faster than lower-level DNN features (see Supplementary Table S4 for more details), and a prediction model using smaller size pooling features converges faster than features with larger pooling size. As a result, a different subset of features may converge to their corresponding optimum at different rates for the same ROI; and the same subset of features may also converge at different rates for different ROIs. Therefore, a single-layer model with a naive concatenating strategy may suffer from the issue of desynchronization for the learned dynamics, and a single model would overfit one ROI and underfit another ROI simultaneously.

To address this issue, we propose that integrating the features across multiple layers with separate optimizations under multiple atlases will improve the prediction accuracy over adopting a single concatenation model.

To test this hypothesis, we implemented the proposed layer-level integration model and compared the model performance against baseline models including concatenation model and single layer encoding models. Specifically, we took a state-of-the-art visual model, the Swin-Transformer model.^44^ We first trained separate encoding models using every intermediate layer of the DNN. These models were optimized end-to-end separately and their backbone Transformer parameters were not fixed. Then we ensembled the outputs of all models through a weighted summation (Fig. 1b), and the ensemble was weighted and optimized using the differential evolution algorithm to maximize the ROI-averaged validation score. This layer-level ensemble model achieved mean *R*^*2*^ = 0.425 on the validation set (Fig. 2a). As a comparison, our full ensemble model dominated the best single-layer model (mean *R*^*2*^ = 0.397) with paired *t*(161325) = 15.7, *p* = 2.21e-55 and the all-layer-concatenation model (mean *R*^*2*^ = 0.376) with paired *t*(161325) = 27.4, *p* = 3.38e-165 under the two-sided two-sample t-test. The significantly improved explained variance of the layer-level integration model over the fully concatenated model indicates the existence of desynchronization in encoding models across layers and suggests the necessity of integrating multi-layer features under various optimization parameters rather than relying on a single model.

**Figure 2.**
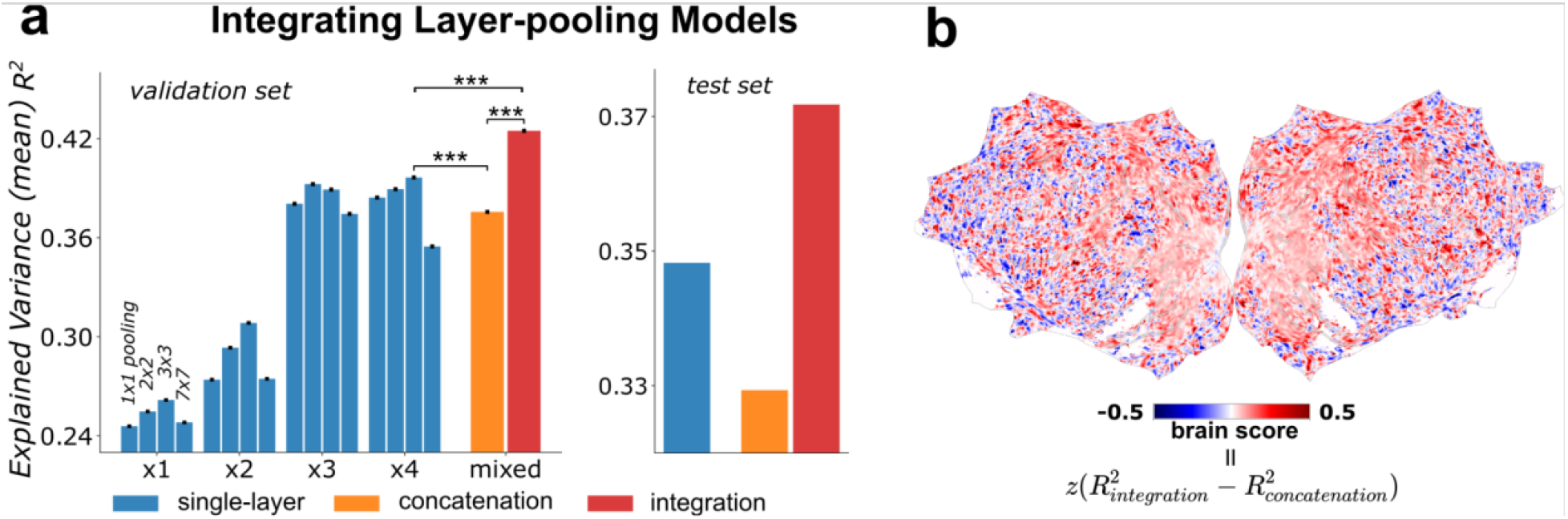
Feature-level integration improves brain prediction performance. a) Averaged brain prediction performance (explained variance) for each individual model. Blue bars: models trained with only one intermediate layer and one pooling size. Orange bar: concatenation model with a naive concatenation of all the input features for blue bar models. Red bar: integrated model that integrates the outputs of blue bar models. b) Cortical mapping of the performance difference between the integration model and the naive concatenation model, scores are noise-normalized.

Furthermore, it is worth pointing out that our model was robust and the results generalized to additional testing sets as well (Fig. 2a). In addition, our Swin-Transformer-based encoding model also outperformed other ensemble models using other architectures, such as 3D ResNet (see Supplement Table S1).

### Constructing hierarchical task-optimized ROI (htROI) atlas

In the previous section, we built a voxel-wise encoding model that integrates DNN representation features across different spatiotemporal scales. The model weights of the encoding model reflected the task-driven functional receptive properties of each individual voxel. To fully exploit the functional structure in the neural activity across the cortex, we next constructed a hierarchical task-optimized atlas (htROI) based on these voxel-wise functional encoding model weights. Specifically, different voxels shared the same parameter in the encoding model except for the last linear layer (Fig. 3a). We concatenated the weights of the last linear layers from multiple models into a vector and used it as the feature vector for each voxel, reflecting task-optimized functional receptive properties.

**Figure 3.**
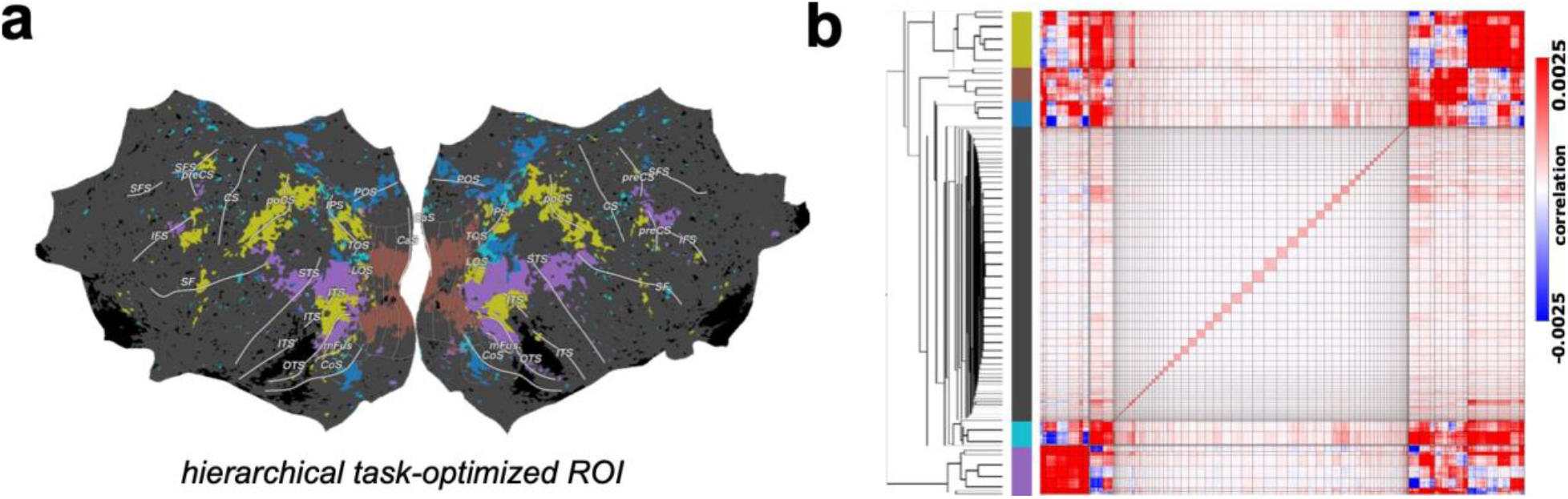
Hierarchical task-optimized ROI (htROI) atlas defined by brain encoding model weights. a) Task-optimized ROI atlas based on hierarchical clustering. Each color represents an ROI, corresponding to the colored column in b). b) Hierarchical clustering: voxels are first clustered by K-means clustering. Vertical and horizontal black lines in the similarity matrix indicate clusters by K-means, each pixel is a voxel pair. An additional hierarchical clustering is performed on K-means cluster centroids, and the final clusters are identified by cutting the dendrogram.

Next, we performed hierarchical clustering^45^ to divide the whole brain into 6 modules (Fig. 3b), including an early visual cluster that mainly covered V1, V2, V3, and V4, a higher-level visual cluster that includes part of the lateral occipital complex (LOC), fusiform gyrus and posterior superior temporal cortex, and a somatosensory cluster that includes the post-central sulcus (Fig. 3a).

### Atlas-level model integration

After building the task-optimized functional atlas, we next integrated voxel-wise encoding models trained on both functional and anatomical atlases to build the final integrated encoding model. Considering the optimization of representation homogeneity within each region, we constructed the prediction model for the voxels in each region separately. We applied the SwinTransformer infrastructure as the backbone and train the prediction model with shared parameters except for the last linear layers. The final voxel-wise neural prediction was a weighted sum of model prediction from all integrated models based on different atlases.

Here we validate whether incorporating brain atlas information into the encoding model would benefit the brain prediction performance, compared to treating whole brain as a homogeneous predicting target. Furthermore, as a functional brain atlas, htROI reflects the functional organization of the voxels, and including htROI in the final integrated model provides additional encoding information that facilitates the brain activity prediction, compared to anatomical-based atlas. To test these hypotheses, we examined our final integrated model performance and compared it against models trained on anatomical atlases only. Specifically, we adopted three atlases that parcellate cortex into different ROIs: the proposed hierarchical task-optimized ROI (htROI), the anatomical ROI (aROI), and the whole-brain ROI (wbROI) that takes the whole brain as a single ROI. The model integrating all three atlases together achieved the best performance on both the validation and test datasets (Fig. 4a, *R*^*2*^ = 0.4686 on the validation set, *R*^*2*^ = 0.3918 on the test set).

**Figure 4.**
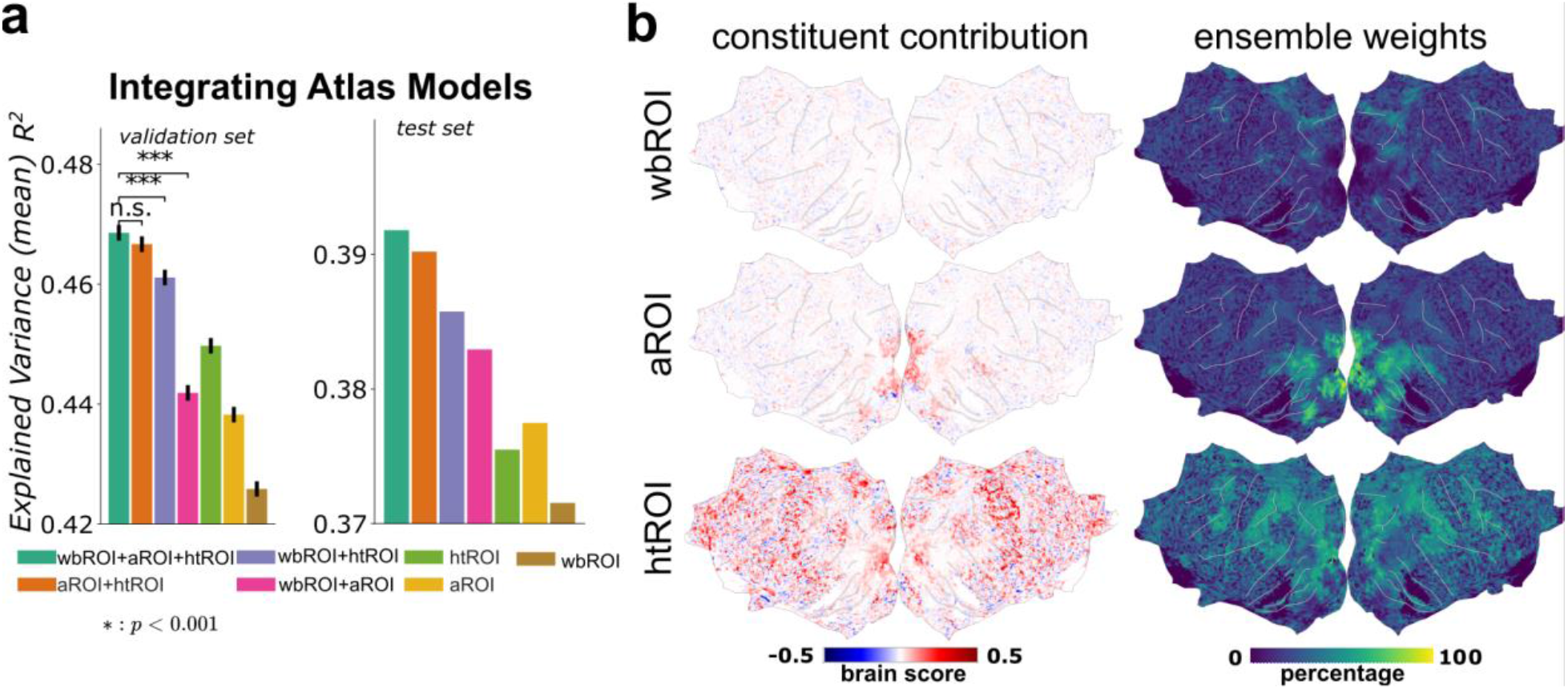
Atlas-level integration further improves brain prediction performance. a) Averaged brain prediction performance over the whole brain (explained variance) for models using different brain atlas partitions (aROI - anatomical ROI partition, htROI - hierarchical task-optimized ROI, wbROI - whole brain). b) Cortical mapping of different atlas-based models. Left panel: constituent contribution measured by the gain in prediction score when adding each atlas model. Right panel: ensemble weight shows the contribution from a specific atlas model in each voxel from a complete ensemble including all atlas models.

**Figure 5:**
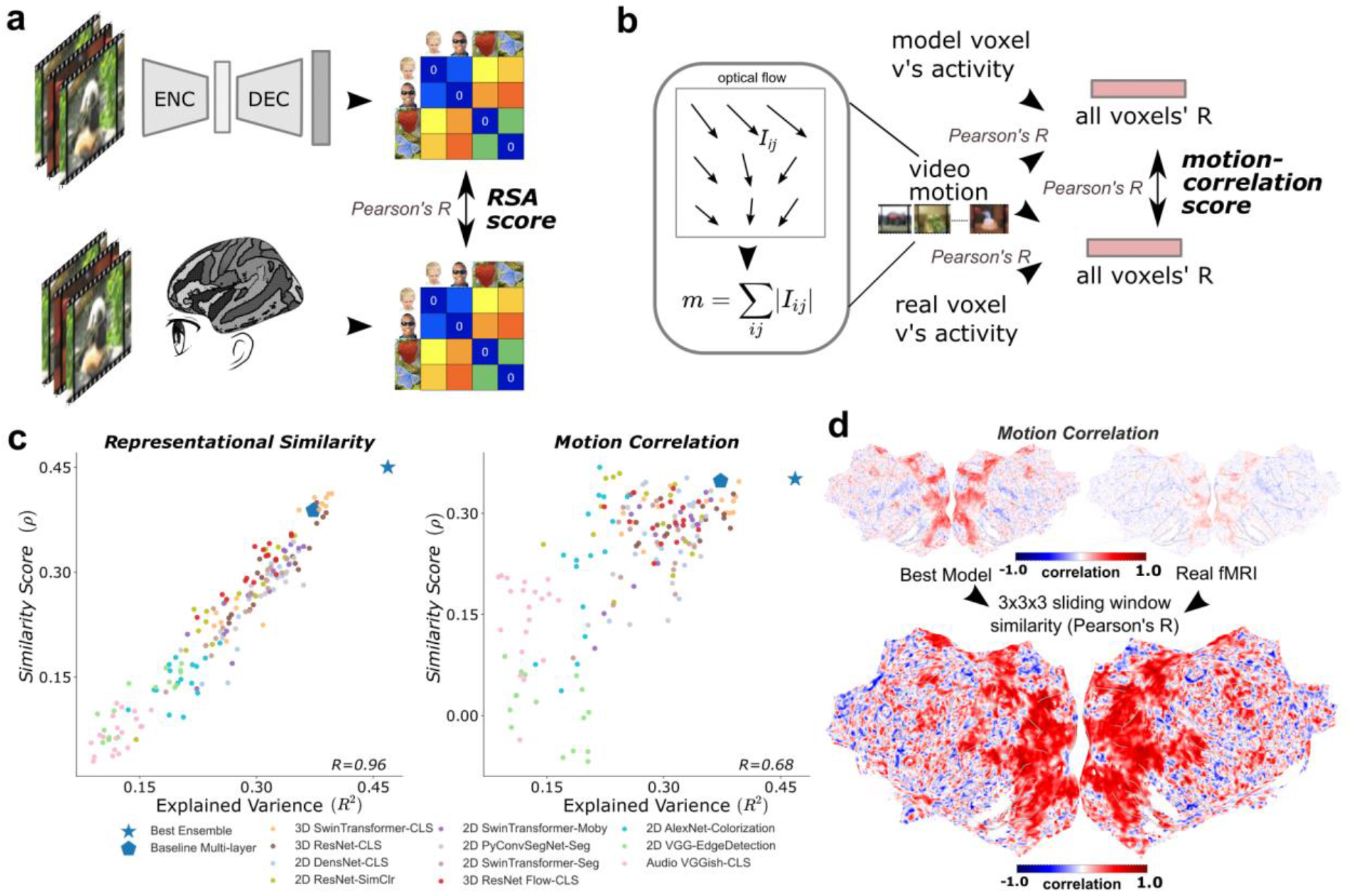
Representation similarity analysis and motion correlation analysis using the proposed integrated encoding model. **a)** Schematic for computing RSA score. We first compute the representation dissimilarity matrix (RDM) in the stimulus space, then compute the similarity score as the Pearson’s correlation coefficient between RDMs from model prediction and from real fMRI signal. **b)** Schematic for computing motion-correlation score. We estimate the motion index as a scalar value for each video by summing all of its optical flow vector magnitudes. The motion-correlation score is calculated by correlating each voxel’s activation to this motion scalar across videos. Finally compare the similarity of motion-correlation score from model prediction and from real fMRI across all voxels. **c)** The correlation between RSA and motion-correlation scores and the brain prediction score of each model (explained variance). Each point is a model with a specific layer-pooling configuration. **d)** *Top*: motion-correlation for each voxel in the integrated prediction model (left) and real fMRI signal (right). *Bottom*: local similarity between the prediction model and real fMRI, estimated as the spatial correlation within the 3 × 3 × 3 sliding window.

To further examine whether the integration is necessary, we performed two levels of ablation study. First, we took the wbROI which obtained *R*^*2*^ = 0.4259 on the validation set and *R*^*2*^ = 0.3715 on test set as the baseline. Both the htROI and aROI outperformed whROI. The aROI obtained *R*^2^=0.4383 and paired *t*(161325) = 6.8, *p* = 1.16e-11 when compared to wbROI under the two-sided two-sample t-test, as well as *R*^*2*^ = 0.3775 on the test set. The htROI obtained *R*^2^=0.4497 and paired *t*(161325) = 13.1, *p* = 3.21e-39 when compared to wbROI under the twosided two-sample t-test, as well as *R*^*2*^ = 0.3755 on the test set. This confirms that incorporating the network module information would contribute to the prediction model. Next, we examined whether the combination of htROI and aROI outperformed each of them separately. The combination of htROI and aROI (i.e., htROI + aROI in Fig. 4a) achieved *R*^*2*^ = 0.4667 and *R*^*2*^ = 0.3902 on the test set. For the comparison, it had paired *t*(161325) = 9.2, *p* =3.26e-20 when compared to htROI and *t*(161325) = 15.4, *p* =1.43e-53 when compared to aROI under the twosided two-sample t-test. This supports the claim that the anatomical and functional atlases contain complementary information to each other and the prediction model benefits from integrating over both atlases. A possible explanation here is that the htROI is designed to maximize the representation similarity in signals of voxels within the same cluster while the aROI provides prior information of module location. Indeed, the improvement of combination over htROI is mainly located on the visual cortex while the improvement over aROI is broadly distributed over the whole brain (Fig. 4b).

### Improvement of conceptual representation through more accurate prediction models

In the previous sections, we built a more accurate model by applying deep neural network models with brain network modularization. The ultimate goal of such models is to better understand neural coding in the brain. Here we demonstrate that with this more accurate voxel-wise prediction model we can better characterize the encoding patterns of image features across the cortex.

Representational geometry of neural populations has been widely studied in neuroimaging to understand the neural coding of sensory information and cognitive processes.^46,47^ Representational similarity analysis (RSA) has become one of the standard methods to compare representations across spaces and to test cognitive and computational theories.^46^ We first analyzed the representation geometry in the predicted activity and the actual BOLD signal using RSA. For each model configuration, we computed the representational dissimilarity matrices (RDMs) of all video stimuli using the model prediction and the actual observed brain responses correspondingly. We then computed a representational similarity score as Pearson’s correlation between the RDMs for the predicted activity of the chosen model and the actual observed brain response. We found that the representational similarity score is strongly correlated with the model’s prediction performance (r = 0.96, p = 3.7e-104) and our proposed model achieved both the highest representational similarity score (*ρ* = 0.4501) as well as the prediction performance (explained variance *R*^2^=0.4686). This indicates that more accurate prediction models also demonstrate more similarity in terms of representational geometry of visual stimuli across the broad visual network in the cortex (Fig. 4).

We next evaluated how our proposed model characterized motion-specific coding in the cortex, which is crucial for analyzing naturalistic video processing. To do this, we defined the motion index in each individual stimulus as the sum of the optical flow vectors’ magnitude. To quantify the neural encoding of motion information, we computed the voxel-wise motion representational similarity, which was Pearson’s correlation between the predicted or actual brain response and the motion index. We found that the prediction accuracy (explained variance) was positively correlated to the motion representational similarity of the predicted neural activity (r = 0.68, p = 1.3e-25), suggesting that our model was able to capture motion-related coding in the brain response. Furthermore, we also evaluated the consistency between the predicted and actual motion representational similarity across the cortex. We found that our model showed high motion coding consistency across a broad range of cortices, including the early visual cortex, the dorsal and ventral visual pathway, and the sensorimotor cortex. This suggests that the performance improvement is beyond simply characterizing low-level texture features in the early visual cortex, but also covers cortical areas involved in intermediate and higher-level information processing.

## Discussion

In this work, we introduced a systematic and data-driven framework of optimizing voxel-wise neural encoding models by integrating DNN representation features and brain network structure information through iterative ensemble learning. Two key ingredients of our proposed method are: 1) the asynchronous integration of multi-scale representation features from DNN models; 2) functional clustering based on encoding model weights, and integration of encoding models over both functional and anatomical atlases. We demonstrated that our proposed method achieved state-of-the-art performance on a large-scale dataset in predicting neural responses to naturalistic videos.

The classical view of visual processing in the cortex supports a domain specific theory of neural coding in the visual cortex with the visual cortex as a hierarchical feedforward processing model.^9,37,48^ These models and theories assume that each cortical area is often exclusively involved in a limited set of functional processing stages and feeds the processed information forward to the next level along the hierarchy. This classical view has guided the computational modeling of the visual cortex in the same way. Previous studies often use a single layer of representation features from pretrained models for a certain ROI.^21,24^ It is also demonstrated that there is a coarse alignment between hierarchical layers in vision CNN and areas in the ventral visual stream.^22^ However, recent studies have challenged this hierarchical idea from anatomical,^37^ experimental^39^ and computational^38^ perspectives, and reveal non-hierarchical processing in the visual cortex. Here we demonstrate a comprehensive framework that exploits the non-hierarchical processing properties by ensembling all different layers of representation from DNN models. Using a data-driven approach, we showed that ensembling lower and higher levels of representations from the DNN hierarchy improved encoding accuracy for both the classical “early” and “late” areas. Our results suggest that both hierarchical and non-hierarchical structures exist in the visual pathway. By evaluating the contributions of different layers and components of the ensemble model, we offer a systematic way of quantifying hierarchical and non-hierarchical structures in the visual system.

The idea of using an *in silico* optimal observer model to infer the underlying computational mechanism in a biological system can be dated back to at least Marr’s three level’s of analysis.^2^ With the emergence of DNN in vision, DNN-based models have been widely adapted as a compositional model of the sensory system, and have shown to be powerful tools in predicting neural activity and behavior.^5^ With a more accurate model, we are able to approach the nonlinear coding property of neural population from a new perspective. Traditional models of single neuron/voxel in the visual system, such as receptive field^49^ or population receptive field models^50^, mostly adopt a theory-driven structural approach. These models mostly use gaussian/gabor filter banks and generalized linear models to denote receptive encoding properties in the image space.^41^ These previous methods are particularly tailored for more intuitive receptive structures in early areas and have been very effective in accounting for important properties, such as retinotopic map. Our approach allows us to evaluate highly abstract, dynamic and nonlinear coding properties in intermediate and higher-level cortical areas,^51^ and account for multi-sensory integration in the more abstract feature embedding space facilitated by the effective ensemble of deep neural network models. These advances allow us to better characterize the neural activity across the cortex.

These more accurate prediction model of the brain can also be used as a preliminary tool to define functional ROI. Our model has shown great ability to generalize across subjects. Thus, we can use such models to define functional ROI based on general naturalistic stimuli without running traditional localizer tasks, which only covers a limited set of stimuli.^52^ This not only saves running time, but also extends the scope of traditional localizer to a novel virtual simulated version. On the other hand, these models also provide novel approach to find optimal stimuli as localizer. Recent study has provided data-driven frameworks to identify optimal stimulus for specific neural circuits using close-loop models.^53,54^ Our model can be fitted into such frameworks and used as the encoder for close-loop brain modulations. In these applications, the ability to accurately predict and generalize to a broad spectrum of input space is crucial.

There are a few aspects that our model can be further improved. Currently we mainly constraint the ensembled models in vision and use the ViT model as the backbone of our specific instantiation of the proposed framework. In a more generalized case, different models from a broader range of modalities can be integrated into the same framework to account for different sensory modalities, such as audition, and to test different hypotheses about neural coding in different cortical networks. Another potential future direction is to explore the generalization and transferability of our proposed approach on different subjects and stimuli as the testing set.

## Methods and Materials

### Dataset in brief

We work on the Algonauts 2021 challenge dataset. Details on data acquisition and preprocessing are provided elsewhere.^40^ Briefly, the dataset consists of 1102 fMRI brain responses per subject (10 subjects), 1000 for training, and 102 held out for online submission. Each stimulus is a 3-second clip of daily events, participants watched the video without playing the sound. Training set videos are scanned 3 times and averaged; test set videos are scanned 10 times to estimate noise ceiling and then averaged. The dataset provides voxel masks for 9 anatomical ROIs (V1, V2, V3, V4, LOC, EBA, FFA, STS, and PPA). BOLD activation is extensively preprocessed by GLMdenoise,^55^ and the stimulus responses are expressed in the regression coefficients of the general linear model. Voxels are filtered by thresholding noise ceiling with 161326 voxels in total for all 10 subjects.

### Voxel-wise encoding model architecture

The voxel-wise encoding model consists of a feature extractor (Video Swin Transformer pre-trained on Something-Something V2 Dataset^56^), a max-pooling read-out head, and a Multi-Layer Perceptron (MLP) prediction head. The outputs is activation values for every voxel in one ROI. One interesting property of this model is that, except for the last linear layer, all the other parameters are shared among all the voxels. This can be formulated as*Y = f*^*1:L-1*^*(X) W*^*L*^, where *Y* ∈ *R*^*B × N*^ is output voxel prediction, B is the batch size, N is number of voxels, *W*^*L*^ ∈ *R*^*D* × *N*^ is the weight for the last layer, D is channel size, *X* is the input video, and *f*^1:*L*−1^ denotes all transformations before the second last layer. We call the parameters in *f*^1:*L*−1^ voxel-shared parameters and *W*^*L*^ voxel-specific parameters (Fig. 1a). *W*^*L*^ contains all the information about an arbitrary voxel, so we use it as task-optimized voxel embeddings for clustering.

### Feature extractor

The deep learning feature extractor model can be formulated as a sequential transformation of input *x*^0^ given by *x*^*L*^ = *δ*^*L*^ ∘ *x*^*L*−1^, where *x*^*L*^is the hidden representation at layer depth *L, δ*^*L*^ is the transformation operation. The pre-trained Video Swin Transformer model consists of 4 major blocks with descending spatial size and increasing channel size (see Table S5 for the details).

### Read-out and Prediction head

We take ***x***^***l***^ and connect it to a read-out head, which consists of an adaptive max-pooling operation with output size ***1*** × ***n*** × ***n, n*** ∈ {***1, 2, 3, 7***}. The output feature of this read-out head is denoted as ***u***^***l***^***n*** = ***pooling***_**1**×***n***×***n***_(***x***^***l***^). The prediction head is a multilayer perceptron (MLP), with Exponential Linear Unit (ELU) activation function on 3 hidden layers, 2048 feature channels per layer. The last layer is set to be without nonlinearity, its output dimension equal to the number of voxels in the ROI.

### Feature-block models ensemble

We train separate models on a cartesian product of all intermediate layers (*l*) and pooling sizes (*n*), *u*^*l*^_*n*_ denotes extracted feature, then ensemble their output *y*^*l*^_*n*_ as described in Algorithm 1. These models are trained to their individual early stopping point. The ensemble is intended to be hierarchical. First, multiple pooling size models are assembled within the same layer, and then ensemble inner-loop outputs are generated. If this hierarchy is violated, the validation score will be overfit and the test score will suffer. (Supplementary Table S1).

##### Algorithm 1. Hierarchical ensemble of separately trained feature-block models

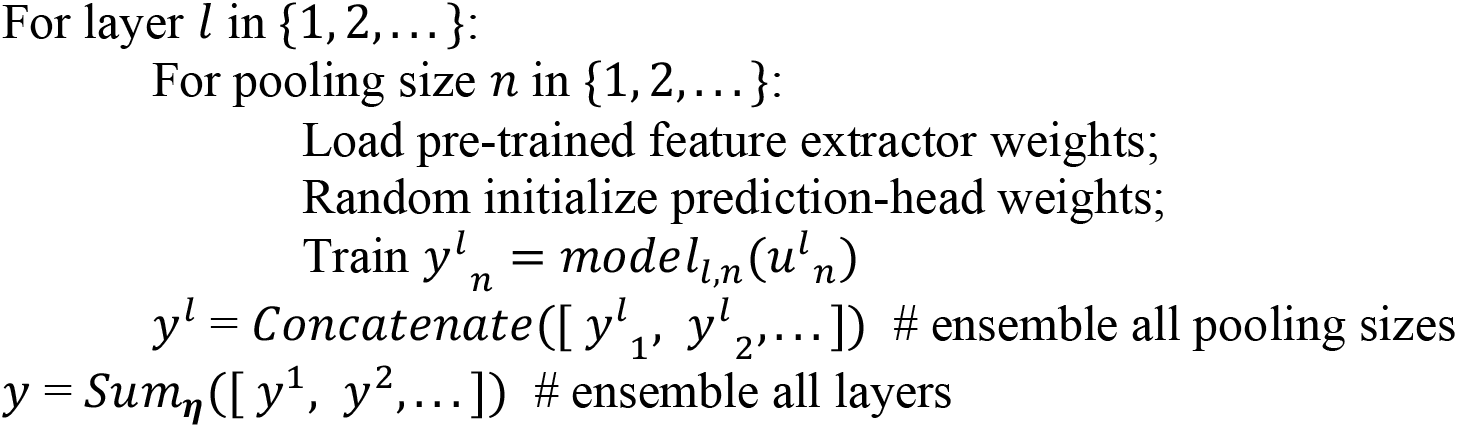

The ensemble operates on model output *y* as

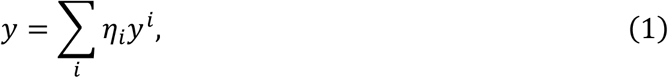

where *η* denote ensemble weights solving *maximize*_*η*_{*σ*(*y*_*val*_, *y*)}, *σ* is the averaged voxel-wise Pearson’s correlation (R) over inputs. The weight *η* is optimized to maximize the prediction score on the validation set through differential evolution.^57^

### Hierarchical task-optimized ROI

We take the last linear layer weight *W*^*L*^ ∈ *R*^*D* × *N*^ as voxel embeddings for clustering. Note that *W*^*L*^ of different models are in separate embedding spaces defined by *f*^1:*L*−1^(*X*), to keep all voxel embeddings in the same space, the model for deriving htROI is trained with all whole-brain voxels. To adapt to ensemble models, we multiply voxel embeddings by their ensemble weight *ηi* and concatenate to obtain a joint voxel embedding *W*^*L*^ = (*η*_1_*W*^*L*^_1_, *η*_2_*W*^*L*^_2_, …, *ηnW*^*L*^*n*), where (*W*^*L*^)^*T*^is then used as input for a K-means (*K* = 100) clustering with euclidean distance, then the cluster centroids are feed to a agglomerative hierarchical clustering with Ward’s method. This 2-stage clustering method help reduce memory and computation usage significantly. Then ROIs are identified by dividing the clustering dendrogram (Fig. S1b Left), note that clusters can be subdivided or merged according to their hierarchy. We also plot a voxel-wise correlation matrix (Pearson’s R) to help identify clusters (Fig. S1b Right).

### Atlases ROI intersection combination

For each voxel v_i, suppose its predicted stimulus induced by the model trained on anatomical atlas is *y*^*s*^_*i*_ and the prediction from the model trained on functional atlas is *y*^*t*^_*i*_, its final output will be a weighted sum from these two models, i.e. *w*^*s*^_*i*_ · *y*^*s*^_*i*_ + *w*^*t*^_*i*_ · *y*^*t*^_*i*_, where *w*^*s*^_*i*_ and *w*^*t*^_*i*_ are the ensembling weight specialized for each voxel. For two or more atlas models, we denote V^*A*^_*i*_ as voxels in the *i* th ROI in atlas A, and V^*B*^ _*j*_ as voxels in the *j* th ROI in atlas B, the intersection of V^*A*^_*i*_ and V^*B*^_*j*_ is V^*AB*^ _*ij*_. Since we trained separate models for each atlas, we ensemble their outputs on V^*AB*^_*ij*_ to maximize prediction score on V^*AB*^_*ij*_. This is repeated for all V^*AB*^_*ij*_ and iterates over all voxels exactly once. The ensemble weight is optimized on the mean of intersection voxels, we also consider voxel-wise ensemble methods, but found voxel-wise methods overfit to the validation set (Supplementary Table S2).

## Data availability

The fMRI data used in this study is available at http://algonauts.csail.mit.edu/challenge.html.

## Code availability

The analysis code used for this study is available at https://github.com/huzeyann/htROI-neural-encoding

## Acknowledgments

The authors gratefully acknowledge the support of the National Natural Science Foundation of China under General Program 61876032 (to S.G.), Shenzhen Science and Technology Program under JCYJ20210324140807019 (to S.G.), Shanghai Pujiang Program under 22PJ1410500 (to Y.L.).

## Author Contributions

Conceptualization: Y.L. and S.G.; Methodology: Y.L., H.Y., and S.G.; Software: Y.L., H.Y., and S.G.; Formal analysis: Y.L., H.Y., and S.G.; Resources: S.G.; Writing: Y.L., H.Y., and S.G.

