## Supplementary Information for "Upgrading Voxel-wise Encoding Model via Integrated Integration over Features and Brain Networks"

Li *et al.*

### Supplementary Information

#### Supplementary Methods.

**Model training.** We first split the 1000 training videos into 90% training and 10% validation. Then, we downsampled all videos to a resolution of $\mathbf{224 \times 224}$ px, $\mathbf{16}$ frames. Subsequent video preprocessing and normalization was identical to that in the pre-trained model. Then we trained the encoding model using the Mean Square Error loss $\frac{\mathbf{1}}{\boldsymbol{n}} \sum_{\boldsymbol{i=1}}^{\boldsymbol{n}} (\boldsymbol{y_i} - \widehat{\boldsymbol{y}_i})^{\mathbf{2}}$, where $\boldsymbol{n}$ denoted the number of voxels in one ROI, $\boldsymbol{y_i}$ was the ground truth voxel activation and $\widehat{\boldsymbol{y}_i}$ was the prediction. At the end of each training epoch, we calculated the correlation between prediction and ground truth on the validation set and averaged it across all voxels. Early stopping was executed if the correlation failed to increase during six consecutive epochs, then restored the model to its best-performing epoch. AdaBelief Optimizer[1] was adopted to train our models, the base learning rate was set to 1e-4, $\boldsymbol{\beta} = (\mathbf{0.9}, \mathbf{0.999})$, $\boldsymbol{\epsilon} = \mathbf{1}\boldsymbol{e} - \mathbf{8}$, weight decouple=True, weight decay=1e-2 for non-bias weights. The training started with the feature extractor model freeze. After reaching a predefined milestone validation correlation score, the backbone is linearly warmed up for 5 epochs starting at 0.1 learning rate ratio. The predefined milestone is set as half of the validation score of a multilayer naive concatenation model with freeze backbone, trained on the same output ROI in a separate debug run. Batch size was set to 32 with gradient accumulating trick to save memory consumption. Pytorch's native support for mixed-precision training (fp16) was employed.

#### Full runtime steps

1. (Notebook000.ipynb) On whole-brain voxels. Launch multiple feature block concatenation baseline model with freeze backbone, record final score and use the 1/2 as backbone defrost milestone.
2. (Notebook001.ipynb) On whole-brain voxels. Launch single layer-pooling feature block models (16 models for each backbone) with backbone defrost milestone score set according to step1.
3. (Notebook010.ipynb) Ensemble all backbone models from step2, also save their voxel embeddings.
4. (Notebook011.ipynb, Notebook012.ipynb) Run hierarchical clustering on voxel embeddings for each backbone model on their weighted and concatenated voxel embeddings. Save the resulting voxels clusters as htROI.
5. (Notebook100.ipynb) On ROI voxels, for each ROI. Launch multiple feature block concatenation baseline model with freeze backbone, record final score and use the 1/2 as backbone defrost milestone.

6. (Notebook101.ipynb) On ROI voxels, for each ROI. Launch single layer-pooling feature block models (16 models for each ROI) with backbone defrost milestone score set according to step5.
7. (Notebook200.ipynb) Ensemble each ROI models from step6. Assemble ROIs from the same atlas to whole-brain.
8. (Notebook200.ipynb) Do ROI intersection ensemble across atlas models.

Table S1. **Comparing different pre-trained backbone models, Video Swin Transformer achieves the best score overall**. Summary of models architecture, input modality, pre-trained dataset, training objective, dimensionality, and prediction brain score. Models are trained with feature-block integration strategy but without atlas ROIs.

| Models | Input modality | Input Resolution | Pre-trained dataset | Unsupervised objective | Supervised objective | # of dimensions in each layer | Brain score (validation/test) |
|---|---|---|---|---|---|---|---|
| Video Swin Transformer[2] | 3D images | 16×224×224 | ImageNet Something-Something v2 | N/A | Classification | 256, 512, 1024, 1024 | 0.4248/ 0.3718 |
| 3D ResNet50[3] | 3D images | 16×224×224 | ImageNet Multi-Moments in Time | N/A | Classification(multi-label) | 256, 512, 1024, 2048 | 0.4246/ 0.3511 |
| DensNet 169[4] | 2+1D images | 16×224×224 | ImageNet | N/A | Classification | 128, 256, 640, 1664 | 0.4230/ 0.3388 |
| ResNet50[5] | 2+1D images | 16×224×224 | ImageNet | Contrastive Learning (SimCLR v2) | N/A | 256, 512, 1024, 2048 | 0.4198/ 0.3330 |
| Swin Transformer[6] | 2+1D images | 16×224×224 | ImageNet | Contrastive Learning (MoCo v2 and BYOL) | N/A | 192, 384, 768, 768 | 0.4021/ 0.3420 |

| | | | | | | | |
|---|---|---|---|---|---|---|---|
| PyConvSegNet[7] | 2+1D images | 16×224×224 | ADE20K | N/A | Semantic Segmentation | 256, 512, 1024, 2048 | 0.4017/ 0.3214 |
| Swin Transformer[8] | 2+1D images | 16×224×224 | ADE20K | N/A | Semantic Segmentation | 128, 256, 512, 1024 | 0.3894/ 0.3253 |
| I3D Flow[9] | 3D images (optical flow) | 64×224×224 | Kinetics400 | N/A | Classification | 192, 480, 832, 1024 | 0.3848/ 0.3215 |
| AlexNet Colorizer[10] | 2+1D images (gray-scale) | 3×56×56 | ImageNet | Colorization | N/A | 128, 512, 512, 313 | 0.3171/ 0.2394 |
| BDCN VGG16[11] | 2+1D images | 3×56×56 | BSDS500 | N/A | Perceptual Edge Detection | 1, 1, 1, 1* | 0.2312/ 0.1887 |
| VGGish[12] | Audio | 3×96×64 | AudioSet | N/A | Classification | 192, 384, 768, 1536, 384 | 0.1888/ 0.1303 |

*: BDCN intermediate layer is scalar pixel edge prediction at different scales.

Table S2. **Integrating intermediate layers: hierarchical Ensemble reduces overfitting to validation score.** Tested on Video SwinTransformer. The hierarchy is defined by Algorithm 1 as nested for-loop, breaking this hierarchy means increasing the parameters in ensemble optimization. Breaking these hierarchies introduce more overfitting as the number of target voxels becomes smaller.

| Ensemble Hierarchy | Atlas(es) | Validation Score (noise-normalized) | Test Score (noise-normalized) |
|---|---|---|---|
| ✓ | wbROI | 0.4259 | 0.3715 |
| | wbROI | **0.4271** | **0.3718** |
| ✓ | aROI | 0.4383 | **0.3775** |

| | | | |
|---|---|---|---|
| | aROI | **0.4397** | 0.3764 |
| ✓ | htROI | 0.4497 | **0.3755** |
| | htROI | **0.4524** | 0.3731 |
| ✓ | wbROI+aROI | 0.4419 | **0.3830** |
| | wbROI+aROI | **0.4439** | 0.3828 |
| ✓ | wbROI+htROI | 0.4611 | **0.3858** |
| | wbROI+htROI | **0.4639** | 0.3846 |
| ✓ | aROI+htROI | 0.4667 | **0.3902** |
| | aROI+htROI | **0.4694** | 0.3885 |
| ✓ | wbROI+aROI+htROI | 0.4685 | **0.3918** |
| | wbROI+aROI+htROI | **0.4714** | 0.3905 |

Table S3: **Integrating atlas models: grouped ensemble on ROI intersection reduces overfitting.** Tested on Video SwinTransformer. Voxel-wise swap to the better model overfit validation score, while intersection swap/ensemble methods reduce this overfitting. The proposed htROI also works significantly better than the aROI (Fig. 4a for htROI vs aROI: $R^2$ = 0.4497 vs 0.4383 mean explained variance for validation set, 0.3755 vs 0.3775 mean explained variance for test set, with paired $t$(161325) = 6.3, $p$ =3.75e-10; for wbROI+htROI vs wbROI+aROI: 0.4611 vs 0.4419 mean explained variance for validation set, 0.3858 vs 0.3830 mean explained variance for test set, with paired $t$(161325) = 10.5, $p$ =1.47e-25).

| Atlas model(s) | Integration strategy | Validation Score (not noise-normalized) | Test Score (noise-normalized) |
|---|---|---|---|
| wbROI+aROI | Voxel-wise swap | 0.5023 | 0.3800 |
| wbROI+aROI | Intersection swap | 0.4374 | 0.3823 |
| wbROI+aROI | Intersection ensemble | 0.4419 | **0.3830** |
| wbROI+htROI | Voxel-wise swap | 0.5424 | 0.3796 |

| | | | |
|---|---|---|---|
| wbROI+htROI | Intersection swap | 0.4517 | 0.3782 |
| wbROI+htROI | Intersection ensemble | 0.4611 | **0.3858** |
| aROI+htROI | Voxel-wise swap | 0.5475 | 0.3827 |
| aROI+htROI | Intersection swap | 0.4567 | 0.3826 |
| aROI+htROI | Intersection ensemble | 0.4667 | **0.3902** |
| wbROI+aROI+htROI | Voxel-wise swap | 0.5778 | 0.3832 |
| wbROI+aROI+htROI | Intersection swap | 0.4582 | 0.3847 |
| wbROI+aROI+htROI | Intersection ensemble | 0.4685 | **0.3918** |

Table S4: **Our motivation for integrations: the desynchronization challenge.** This table shows the training epochs until reaching the best validation score, early stopping is configured on the average score of voxels from one ROI. Concatenating layers and pooling sizes lead to the lowest training time compared to separate models, which implies it converged to the closest local minimum. Combining output voxels (WB vs V1/STS) leads to a different distribution of training time, which implies it is not the optimal point for each individual ROI. WB: whole-brain voxels.

| Optimal Epochs (V1, STS, WB) | | Pooling Size(s) | | | | |
|---|---|---|---|---|---|---|
| | | 1×1 | 2×2 | 3×3 | 7×7 | ALL |
| Layer(s) | x1 | 12, 17, 24 | 15, 18, 30 | 13, 20, 31 | 14, 12, 23 | \ |
| | x2 | 18, 11, 22 | 16, 12, 26 | 13, 11, 27 | 10, 11, 18 | \ |
| | x3 | 17, 16, 28 | 19, 15, 28 | 13, 12, 27 | 8, 9, 20 | \ |
| | x4 | 18, 14, 31 | 14, 13, 28 | 13, 17, 28 | 9, 8, 22 | \ |
| | ALL | \ | \ | \ | \ | 8, 8, 19 |

Table S5. **Feature extractor dimensions.**

| Depth | $x^0$ | $x^1$ | $x^2$ | $x^3$ | $x^4$ |
|---|---|---|---|---|---|
| Dimensions | 16×224×224×3 | 8×56×56×256 | 8×28×28×512 | 8×14×14×1024 | 8×7×7×1024 |

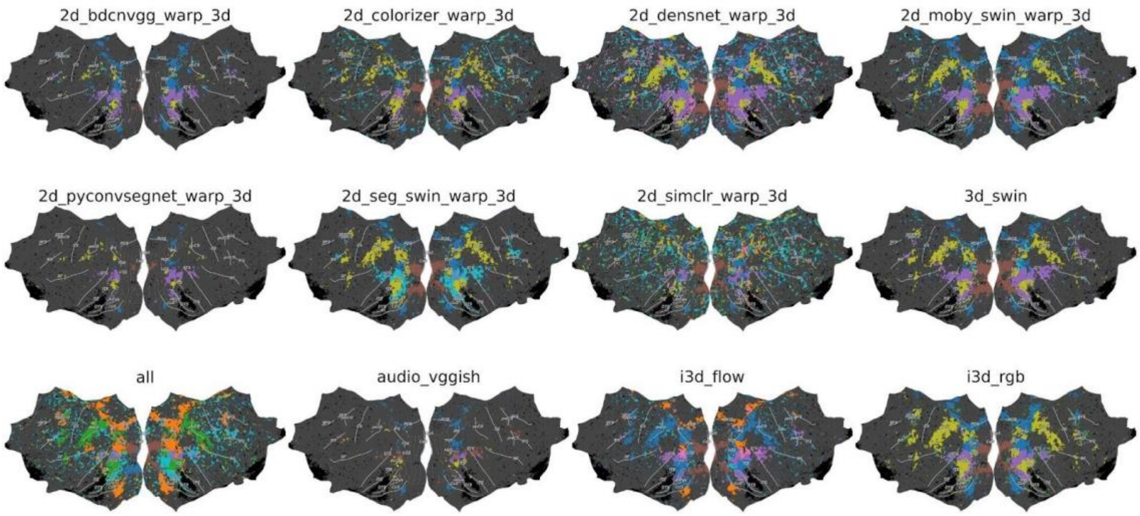


**Figure S1: The htROIs derived from all types of models in Table S1.**

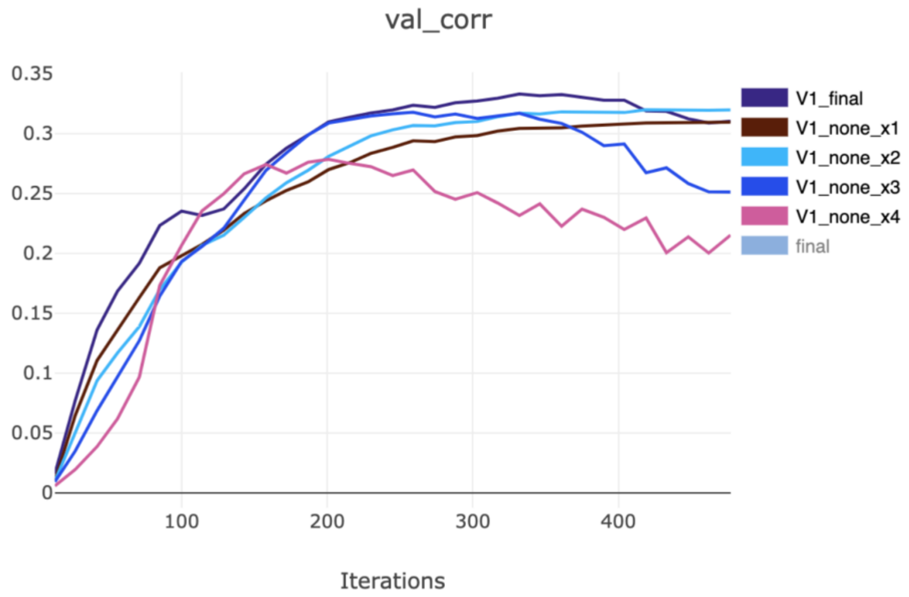


**Figure S2: The desynchronization problem of intermediate layers.** Here the problem is that $\boldsymbol{x_4}$ and $\boldsymbol{x_1}$ converge at different speeds. Thus it is difficult to synchronize the training for them simultaneously. If we enforce them to adopt the same ending epoch, the final score would be lower than each own best layer score.
